# Effect of Handedness on Learned Controllers and Sensorimotor Noise During Trajectory-Tracking

**DOI:** 10.1101/2020.08.01.232454

**Authors:** Momona Yamagami, Lauren N. Peterson, Darrin Howell, Eatai Roth, Samuel A. Burden

## Abstract

In human-in-the-loop control systems, operators can learn to manually control dynamic machines with either hand using a combination of reactive (feedback) and predictive (feed-forward) control. This paper studies the effect of handedness on learned controllers and performance during a trajectory-tracking task. In an experiment with 18 participants, subjects perform an assay of unimanual trajectory-tracking and disturbance-rejection tasks through second-order machine dynamics, first with one hand then the other. To assess how hand preference (or dominance) affects learned controllers, we extend, validate, and apply a non-parametric modeling method to estimate the concurrent feedback and feedforward controllers. We find that performance improves because feedback adapts, regardless of the hand used. We do not detect statistically significant differences in performance or learned controllers between hands. Adaptation to reject disturbances arising exogenously (i.e. applied by the experimenter) and endogenously (i.e. generated by sensorimotor noise) explains observed performance improvements.

## I. Introduction

HUMANS interact with dynamic machines and devices such as computers, quadrotors, and cars in daily life. These interactions give rise to a *human-in-the-loop* control system where the human and the machine jointly accomplish a task through one or more sensorimotor loops. For instance, in trajectory-tracking tasks, people can visually observe the machine and provide input through a manual interface like a mouse, joystick, or steering wheel [1]–[9]. In such cases, people learn to steer computer cursors, quadrotor drones, and personal vehicles using *visuomotor control*. Such manual interfaces often prescribe *how* we interact with the system: some tasks are performed with one hand, others require coordination between hands, and still others may use either or both left and right hands (e.g. the mouse, joystick, and steering wheel, respectively). Because performance in tasks involving fine motor control is affected by the hand used [10], we seek to understand how human visuomotor control differs between hands toward developing effective human-in-the-loop systems. For instance, modeling differences in control between hands could be used to improve bimanual interfaces or to assist unimanual interaction when someone’s preferred hand is unavailable due to injury, disease, or circumstance.

Colloquially understood as the “differences between the hands in terms of skill” [10], handedness can be quantitatively assessed with questionnaires (e.g. the Edinburgh Handedness Inventory [11] or Annett Handedness Questionnaire [12]) or observed from dexterity tasks [12] when questionnaires are difficult or unreliable to administer (such as for young children). These assessments suggest that about 63% prefer to use the right hand and about 7% prefer to use the left hand [12]. This means that about 70% of people have a *preferred* or *dominant* hand that is more dexterous than the *non-preferred* or *non-dominant* hand. Ongoing research indicates that the observed differences in dexterity between dominant and non-dominant hands may be due to each hemisphere of the brain specializing for different aspects of limb movements (termed *lateralization*) [13]–[15].

Studies in sensorimotor neuroscience suggest that participants learn different sensorimotor skills with their dominant versus non-dominant hand. For instance, when performing a reaching task under the influence of a force field applied by a robotic manipulandum, participants learned to improve final position accuracy for both dominant and non-dominant hands [13]. However, initial movement direction improved only for the participants’ dominant hand, which the researchers attribute to changes in predictive (i.e. feedforward) control, whereas the non-dominant hand primarily improved in final error correction, which the researchers attribute to changes in reactive (i.e. feedback) control. These findings suggest that participants rely more on feedforward than feedback control when using their dominant hand, and vice-versa when using their non-dominant hand, for reaching tasks [13], [15]–[18].

For continuous trajectory-tracking and disturbance-rejection tasks through (smooth non)linear machines, prior research primarily focused on modeling participants using their dominant hand [1]–[8], [19]. The results from these experiments support the hypothesis that humans learn to use a combination of feedback and feedforward control to reject disturbances and track references. However, little is known about the differences between controllers learned with different hands and whether learned controllers transfer between hands [9]. We seek to determine whether the differences in control mechanisms between left and right hands found in rapid reaching tasks [13], [15]–[17] extend to continuous trajectory-tracking tasks.

The goal of this paper is to determine whether participants learn different feedback or feedforward controllers when using their dominant versus non-dominant hand during a visuomotor trajectory-tracking task. We extend, validate, and apply a non-parametric system identification method to estimate feedback and feedforward controllers using unpredictable reference and disturbance signals and second-order machine dynamics. Then we experimentally assess differences in sensorimotor learning between the dominant and non-dominant hand and test whether controllers transfer between hands.

We previously reported preliminary results for first-order machine dynamics in a non-archival conference proceeding [5]; this paper extends those results to a second-order system and provides additional support for the underlying assumptions and hypotheses. More significantly, this paper presents new results comparing learned controllers and performance obtained with dominant and non-dominant hands.

Specifically, two groups learned to perform a unimanual trajectory-tracking and disturbance-rejection task. One group started with their dominant right hand before switching to their non-dominant left hand, and vice-versa for the other group. To assess the effect of handedness on learning and transfer, we compared (i) feedback and feedforward controllers and (ii) performance obtained by the two groups with their dominant and non-dominant hands. We found that handedness did not affect the learned controller during a continuous trajectory-tracking and disturbance-rejection task. Additionally, we provide evidence that improvements in trajectory-tracking performance may be attributed to changes in feedback gain to reject disturbances applied (a) externally by the experimenter, leading to system-level performance improvements only for the group that learned the task with their non-dominant hand first, and (b) internally due to sensorimotor noise.

## II. Background

We adopt a tutorial expository style in this section for two reasons. First, to support validation of the assumptions underlying our modeling and analysis methodology, it is important that we explicitly state these assumptions. Second, to support the application of our methods outside the human-in-the-loop controls community, it is valuable to explicitly provide details and rationale that would ordinarily be taken as common knowledge in our niche community. The expert reader may wish to skim or skip this section after reviewing the following table of symbols and Fig. 1, returning only if questions arise in subsequent sections.

**Fig. 1:**
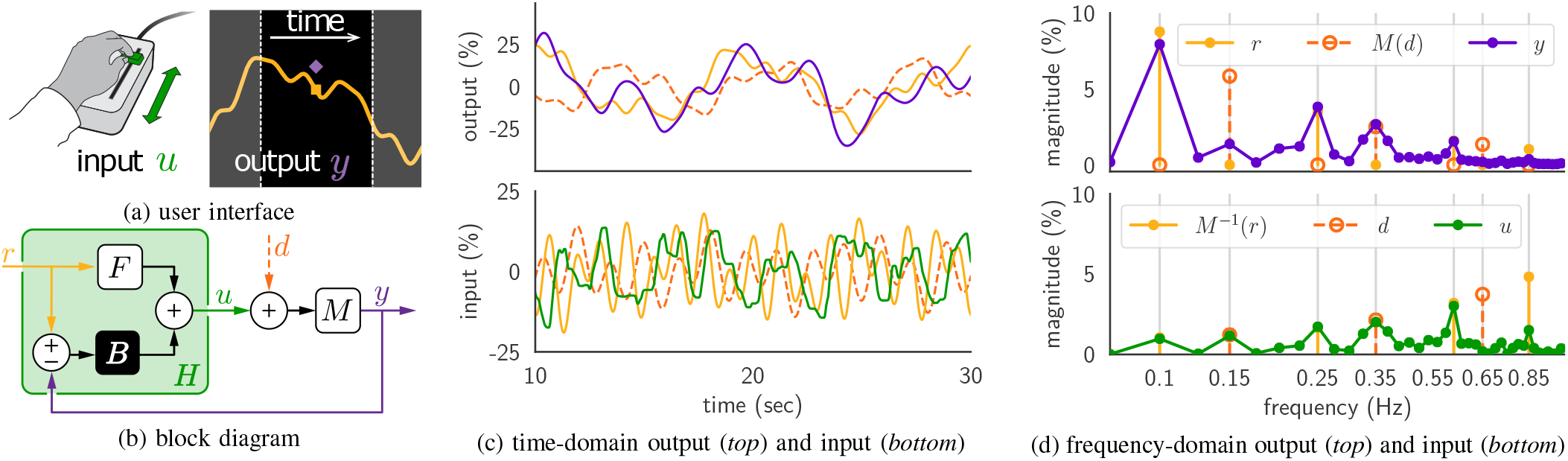
Human-in-the-loop trajectory-tracking. (a) Human response 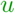 is obtained with a one-dimensional manual slider and input to machine *M* to produce output 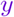, which is overlayed on a display with 1 sec of a reference trajectory (0.5 sec preview). (b) The human 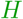 transforms reference 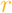 and output 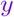 to user response 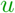; the machine *M* transforms the sum of control 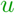 and disturbance 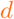 to output 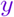. We hypothesize that the human’s transformation is the superposition of a *feedforward F* response to reference 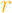 and a *feedback B* response to tracking error 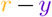. Representative data from one trial of the **linearity** experiment are shown in (c) the time-domain and (d) the frequency-domain. The frequency content of 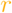 and 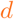 are confined to prime multiples of a base frequency (1/20 Hz). Magnitudes shown as percent of output or input space extent.

### A. Response to Reference and Disturbance Superimposes

In the laboratory, we instantiate the human-in-the-loop system as a one-degree-of-freedom reference-tracking and disturbance-rejection task (Fig. 1) [2]. The transformations that must take place inside the human (i.e. to observe cursor position, generate a motor plan, and control muscles to move the hand) are known to be nonlinear. However, when tasked with tracking reference *r* and rejecting additive disturbance *d* through a linear time-invariant (LTI) [20, Ch. 3, pg. 4] system *M*, we assume that people behave approximately like LTI transformations for a range of reference and disturbance signals [2], [4]–[6], [9], [21]. When this assumption holds, the control signal *u* produced by the human in response to reference *r* and disturbance *d* satisfies the law of superposition,

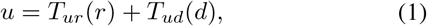

where *T_ur_* and *T_ud_* are LTI transformations.

#### Hypothesis 1.

*The user response for reference tracking with disturbance is consistent with a superposition of the user response to the reference and disturbance signals presented individually.*

Signals and LTI systems have time-domain and frequency-domain representations as in Fig. 1(c,d), related by the *Fourier transform* [22, Ch. 5]; we will adorn signal *x* and transformation *T* with a “hat” 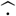 to denote the Fourier transform 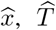. Importantly in what follows, the frequency-domain operation performed by an LTI system is particularly simple: each frequency component of the input is independently scaled and phase-shifted [20, Ch. 9]. Thus, frequency-domain LTI transformations (termed *transfer functions*) can be empirically estimated by dividing Fourier transforms of time-domain input and output signals at each frequency of interest *ω* and visualized using a *Bode plot* [22, Ch. 5] as in Fig. 4. Specifically, when disturbance *d* = 0 in (1) we have

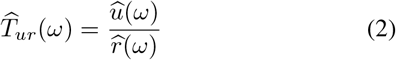

and when reference *r* = 0 in (1) we have

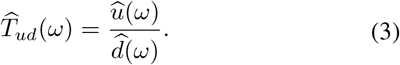

In contrast, an LTI system’s time-domain operation (1) – *convolution* [22, Ch. 3] – is mathematically and computationally more complicated than frequency-domain multiplication. For this reason, we design and analyze experiments using frequency-domain representations of signals and systems.

### B. Combined Feedback and Feedforward Improves Prediction

In the absence of reference, (i.e. *r* = 0), we assume that the human response is solely due to a *feedback B* transformation of *tracking error e* = *r* − *y* = −*y* (i.e. *H*(0, *y*) = *B*(−*y*)). If a nonzero reference *r* ≠ 0 is known to the human, we assume that it evokes an additive *feedforward F* transformation of *r*, so that the overall human response can be written as

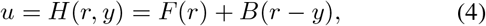

where *e* = *r* − *y* is tracking error. Using a combination of feedback and feedforward control to model human trajectory-tracking has a long history in the field [1]–[3], [5]–[7], [9], [23]–[25], and is a well-known strategy to improve performance over error feedback alone [20, Ch 8]. We emphasize, however, that certain neurologic conditions like cerebellar ataxia could impair people’s ability to perform feedforward control. In such cases, feedback alone may provide better predictions [26], [27].

Under Hypothesis 1, we can apply *block diagram algebra* [20, Sec. 2.2] to transcribe Fig. 1(b) into equations that can be manipulated to express the empirical and prescribed transfer functions 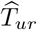 (2), 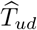 (3), 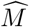 (7) in terms of the unknown transformations *F* and *B*,

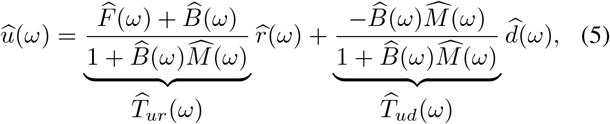

and solve (5) to estimate the feedback *B* and feedforward *F* components of the human’s controller at each stimulated frequency *ω*,

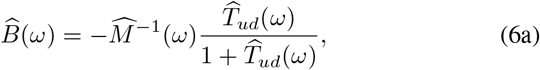

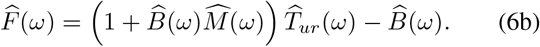

If we instead, assume that the human’s response to reference *r* is entirely due to feedback *B*, then the feedforward controller *F* estimated in (6b) will be approximately zero, so it can be neglected in (4) without affecting prediction accuracy.

#### Hypothesis 2.

*The combined feedback and feedforward model predicts user responses better than a solely feedback model.*

### c. Feedback and Feedforward Adapt with Experience

Previous studies on point-to-point reaching tasks suggest that improvements in end-point accuracy can be attributed to improvements in initial movement (feedforward control) for the dominant hand and improvements in error correction (feedback control) for the non-dominant hand [13], [16], [17], possibly due to specialization of each arm and the corresponding brain hemisphere that controls the arm [13]–[15]. These findings lead to the hypothesis that similar observations will hold in the trajectory-tracking task considered here.

#### Hypothesis 3.

*Human feedback and feedforward controllers will adapt with practice. (a) Feedback will adapt when using the non-dominant hand. (b) Feedforward will adapt when using the dominant hand.*

Note that this hypothesis does not speculate about *how* controllers adapt in the trajectory-tracking task.

## III. Experimental Methods

Two experiments approved by the University of Washington, Seattle’s Institutional Review Board (IRB #00000909) were conducted to:

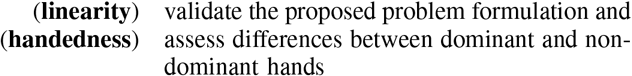

during sensorimotor learning and control in a continuous trajectory-tracking task.

### A. Manual Interface

Participants used a one-degree-of-freedom manual interface to control the position of a cursor on a screen to track a reference trajectory (Fig. 1a). The interface handle was attached to a linear potentiometer; the user response *u* was determined by measuring the potentiometer voltage using an Arduino Due (`Arduino.cc`). The linear potentiometer had a 10 cm extent, and trials were designed such that the input required to produce the reference trajectory was restricted to the middle third of this physical extent. The handle geometry changed between the **linearity** and **handedness** experiments to improve ergonomics (Fig. 2):

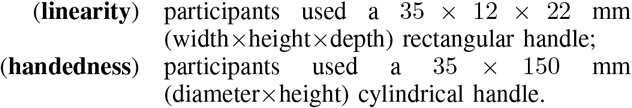

**Fig. 2:**
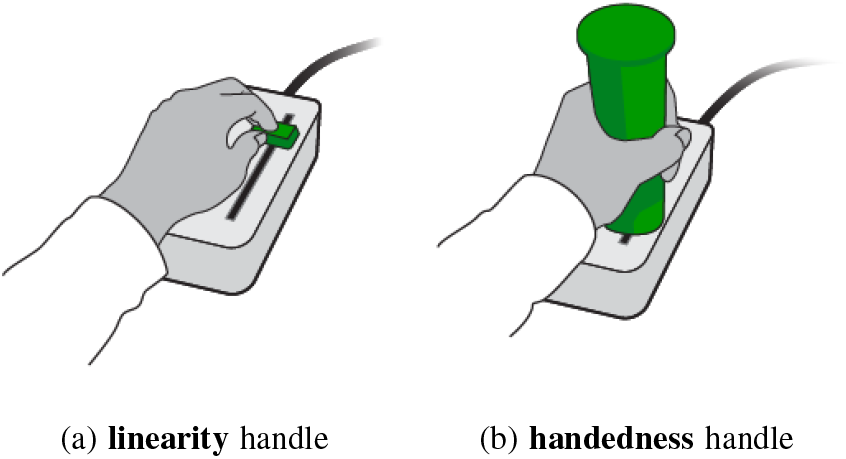
Handles for **linearity** and **handedness** experiments. (a) Participants pinched a rectangular handle with their fingers in the **linearity** experiments. (b) Participants grasped a cylindrical handle with their hand in the **handedness** experiments.

### B. Unpredictable Stimuli

Reference and disturbance signals were constructed as a sum of sinusoidal signals with distinct frequencies. Each frequency component’s magnitude was normalized by the frequency squared to ensure constant signal power, and the phase of each frequency component was randomized in each trial to produce pseudorandom time-domain signals as in Fig. 1(c). A similar stimulus design procedure was employed in [6] to produce unpredictable reference and disturbance signals, and in [1] to produce unpredictable disturbance signals. However, to prevent harmonics from confounding user responses at different frequencies, we adopted the procedure from [28] that restricts stimuli frequency components to prime multiples of a base frequency (1/20 Hz in our experiments). Each trial consisted of two periods of the periodic stimuli (40 sec total) after a 5 sec ramp-up. The number of prime multiples changed between the **linearity** and **handedness** experiments to balance the experiment design:

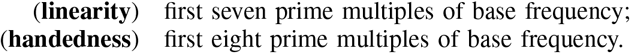

### C. Trajectory-Tracking Task

User response *u* was transformed through a second-order system with damping to produce output *y*:

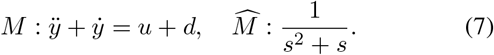

In all experiments, 1 second of reference *r* was displayed with 0.5 second preview, participants were tasked with adjusting their control *u* to make a cursor positioned at *y* track the reference, and the user’s response *u* was modified by an additive disturbance *d* to determine the machine output *y* = *M* (*u* + *d*).

#### 1) Conditions for Linearity Experiment

To test the super-position principle (1), the three different types of conditions illustrated in Fig. 3 were presented to the user in the order shown in TABLE II. In disturbance-only trials (condition (0, *d*)), the reference *r* was constant (zero) and the disturbance *d* was non-constant. In reference-only trials (condition (*r,* 0)), the reference *r* was non-constant and the disturbance *d* was zero. In reference-plus-disturbance trials (condition (*r, d*)), both signals were non-constant, but their frequency components were interleaved as in Fig. 3 (*bottom*) to distinguish the user’s response to both signals: specifically, reference or disturbance were active at even or odd multiples of the base frequency (indicated by an *E* or *O* subscript, respectively). The two types of (*r, d*) trials – (*r_E_, d_O_*) and (*r_O_, d_E_*) – were presented to the participants in alternating order.

**Fig. 3:**
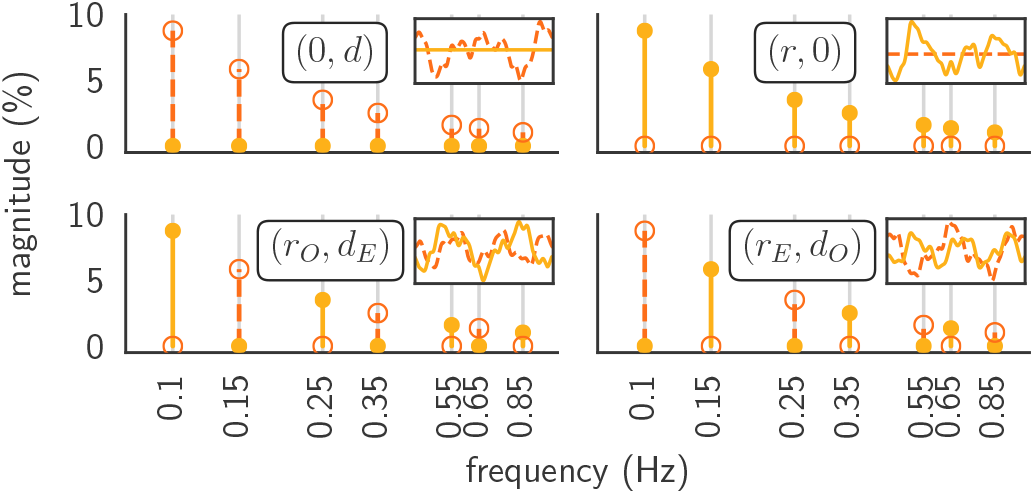
Conditions for **linearity** experiment (cf. TABLE II). To assess whether the human’s response to external reference *r* superimposes with the response to external disturbance *d*, we empirically estimated transfer functions using data from four experimental conditions: disturbance-only ((0, *d*), upper left); reference-only ((*r,* 0), upper right); reference and disturbance interleaved at different frequencies ((*r, d*), bottom left, right). The magnitude of 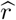 is denoted with solid lines and filled circles, while dashed lines and open circles denote that of 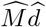; insets show corresponding time-domain signals *r*, *M* (*d*). Magnitudes shown as percent of output or input space extent.

**TABLE I:** Table of symbols.

| symbol | reference | meaning |
| --- | --- | --- |
| $u$ | Fig. 1 | human response signal |
| $y$ | Fig. 1 | machine output signal |
| $r$ | Fig. 1 | reference trajectory signal |
| $d$ | Fig. 1 | input disturbance signal |
| $T_{zx}$ | Sec. II-A | transformation from signal $x$ to signal $z$ |
| $T(x)$ | Sec. II-A | transformation of $x$ by $T$ |
| $\hat{x}, \hat{T}$ | Sec. II-A | Fourier transformation of $x, T$ |
| $M$ | Sec. II-B | machine transformation: $y = M(u + d)$ |
| $H$ | Sec. II-B | human transformation: $u = H(r, y)$ |
| $B$ | Sec. II-B | human feedback controller |
| $F$ | Sec. II-B | human feedforward controller |

**TABLE II:** Conditions for **linearity** experiment (cf. Fig. 3).

| Order | 1 | 2 | 3 | 4 | 5 |
| --- | --- | --- | --- | --- | --- |
| Condition | $(r, d)$ | $(0, d)$ | $(r, d)$ | $(r, 0)$ | $(r, d)$ |
| # Trials | 2 | 10 | 2 | 10 | 10 |

#### 2) Conditions for Handedness Experiment

To assess the effects of handedness on feedback and feedforward control, participants were divided into two groups. All participants were right-handed, so we refer to the dominant hand as the “right” hand and the non-dominant hand as the “left” hand. The first group completed 30 (*r, d*) trials with their dominant right hand, then 30 (*r, d*) trials with their non-dominant left hand (Group RL). The second group completed the same number of trials, but with their non-dominant left hand first, followed by their dominant right hand (Group LR).

### D. Data Analyses

User response *u*, reference *r*, disturbance *d*, and output *y* were sampled at 60 Hz and converted to frequency-domain representations using the fast Fourier transform (FFT). Data were analyzed using `Python3.5`. Transfer functions were estimated at stimulated frequencies from distributions obtained using equations (2), (3), and (6); this simple non-parametric modeling scheme is referred to as the *Fourier coefficients method* [29] or the *spectral measurement technique* [6].

#### 1) Hypothesis 1

We computed frequency-domain representations of the transformation from disturbance *d* and reference *r* to response *u* (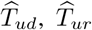 respectively) at each stimulated frequency using (2) and (3). We performed a Wilcoxon signed-rank test with significance threshold *α* = 0.05 to assess whether the magnitudes and phases of 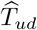 and 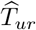 in the (0, *d*) and (*r,* 0) trials were different from those in the (*r, d*) trials. The Wilcoxon signed-rank test is a non-parametric paired *t*-test for data that is not normally distributed [30, Sec. 5.7], selected for this study due to the small expected sample size of less than 10 participants (see Appendix C for more details). If there are statistically significant differences between transformations estimated from different conditions, it suggests that the human is not well-modeled as an LTI system.

#### 2) Hypothesis 2

Feedback *B* was estimated for each participant by applying (6a) to data from disturbance-only trials (condition (0, *d*)) and averaging across trials; similarly, feed-forward *F* was estimated for each participant by applying (6b) to data from reference-only trials (condition (*r,* 0)), using *B* that was just estimated from the (0, *d*) trials and averaging across trials. These controller estimates were used to predict user response 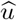 by applying (5) to data from disturbance-plus-reference trials (condition (*r, d*)) for the last 10 trials. The coefficient of determination *R*^2^ [31, Eqn. (3.9)] was used to assess prediction accuracy at each frequency (see Appendix A for more details). We assessed differences between the *R*^2^ value obtained from the feedback-only (*B*) model and the feedback-plus-feedforward (*B* + *F*) model with the Wilcoxon signed-rank test with significance threshold *α* = 0.05. If there is a statistically significant improvement in the *R*^2^ value for the *B* + *F* model compared to the *B*-only model, it suggests that the human response is better modeled with a combination of feedback and feedforward control.

#### 3) Hypothesis 3

We assessed the performance of each participant using time-domain tracking error computed as the mean-square error (MSE) between reference *r* and output *y*:

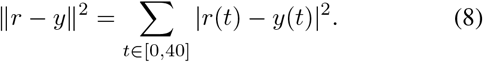

Changes in performance over time were assessed by applying the Wilcoxon signed-rank test with *α* = 0.05 to the average performance of each individual over the first and last five trials with each hand. To assess differences between Group RL and Group LR, we performed the Mann-Whitney *U* test, a non-parametric unpaired *t*-test, with *α* = 0.05.

To assess whether a transformation *T* changed with practice, we averaged the magnitude of the frequency-domain representation 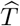 at stimulated frequencies *ω* ∈ {0.10 Hz, 0.15 Hz},

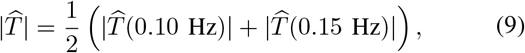

averaged this quantity over the first and last five trials with each hand for each participant, and applied the Wilcoxon signed-rank test with *α* = 0.05. We only included the first two stimulated frequencies in (9) since the other stimulated frequencies exceeded the crossover frequency^1^ observed in our population, and prior work indicates (and our results corroborate) that reference-tracking and disturbance-rejection performance degrades at frequencies higher than crossover.

This procedure was applied to the estimated human feed-forward 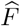 and feedback 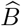 transformations, as well as the system-level transformations 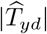 and 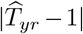. Our focus on the latter two transformations is motivated by the observations that the disturbance is rejected if 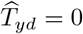 and the reference is tracked if 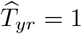. However, we note that 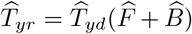 (assuming *F* and *B* are LTI), so it is not possible for the user to simultaneously achieve 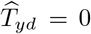 and 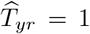 (assuming 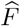 and 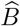 have finite magnitude). We quantify *system-level performance* at each frequency using statistics for trajectory tracking and disturbance rejection, 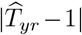 and 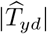. Smaller values correspond to better performance.

#### 4) Non-Stimulated Frequencies

Our methods can only estimate transfer functions at stimulated frequencies; the denominators in (2) and (3) are undefined at frequencies *ω* where 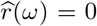 or 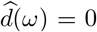, respectively. Although we expect the power of the user response signal to be concentrated at these stimulated frequencies, we nevertheless measure user response at intermediate non-stimulated frequencies (see Fig. 1(d)). Since any user response at non-stimulated frequencies degrades task performance (there is no reference to track or disturbance to reject), we use the deviation of 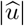 from 0 as another way to quantify task performance. Tracking error (8) is affected by user response across both stimulated and non-stimulated frequencies, which we separately quantify using system-level performance and 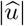.

## IV. Results

We recruited participants from the greater University of Washington community: 7 for the **linearity** experiment, and an additional 18 (9 male, 9 female; age 18-32; height 145-190 cm; weight 48-98 kg) for the **handedness** experiment.^2^ The participants had no reported neurological or motor impairments and all were daily computer users.

### A. Response to Reference and Disturbance (Approximately) Superimposed Across Conditions

We tested Hypothesis 1 with the **linearity** experiment to determine whether user response *u* in disturbance-only (0, *d*) or reference-only (*r,* 0) conditions was consistent with user response in disturbance-plus-reference conditions (*r, d*) (Fig. 4). The magnitude and phase of the transfer functions from *d* and *r* to *u* (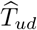 and 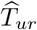, respectively) estimated from these different conditions were indistinguishable at most stimulated frequencies (*p* > 0.05; exceptions denoted with † in Fig. 4), indicating that participants’ response to reference and disturbance signals approximately satisfied the law of superposition across the qualitatively different conditions in Fig. 3.

**Fig. 4:**
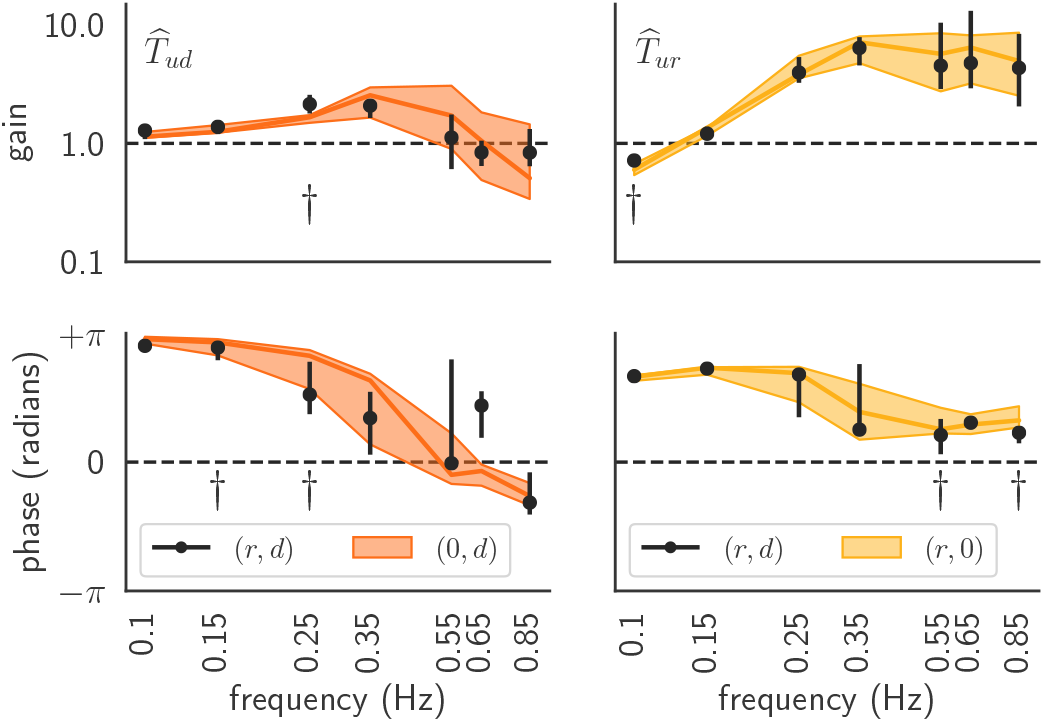
Transfer function estimates in **linearity** experiment. Distributions (median, interquartile) of transfer functions 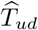 (*left*), 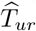 (*right*) estimated from disturbance-only or reference-only trials, (0, *d*) or (*r,* 0), and reference-plus-disturbance trials (*r, d*), for the conditions in TABLE II and Fig. 3. Statistically significant differences (Wilcoxon signed-rank test: *p* < 0.05) in distribution magnitude or phase at each frequency indicated with †.

### B. Combined Feedback and Feedforward Improved Prediction

We tested Hypothesis 2 with the **linearity** experiment to determine whether a combined feedback-plus-feedforward (*B* + *F*) model improves prediction compared to a feedback-only (*B*) model (Fig. 5). Predictions for both models were better (*R*^2^ closer to 1) below crossover frequency (0.25 Hz, determined as the lowest stimulated frequency where the gain of the open-loop transfer function 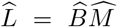 is less than 1 [2], [6]), and decreased in accuracy (*R*^2^ closer to 0) at higher frequencies, suggesting the linear models were more accurate at lower frequencies. Prediction accuracy for the *B* + *F* model was higher than the *B* model at all frequencies (*Z* = 0.0, *p* = 0.016), suggesting that user responses *u* to references *r* and disturbances *d* are better predicted with a combined feedback-plus-feedforward (*B* + *F*) model than a feedback-only (*B*) model.

**Fig. 5:**
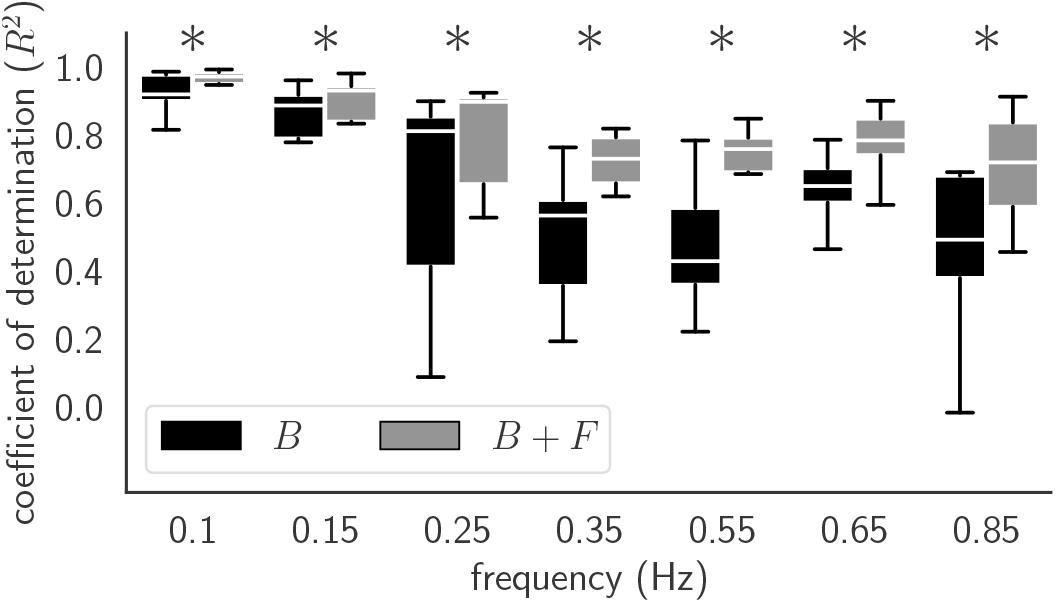
Predictive accuracy of models, **linearity** experiment. Distribution (median, interquartile, range) of coefficient of determination (*R*^2^) between human inputs *u* and predictions from feedback-only (*B*) and feedback-plus-feedforward (*B* + *F*) models. The *B* + *F* model had significantly better prediction accuracy than the *B* model at all frequencies (Wilcoxon signed-rank test: *Z* = 0.0, *p* = 0.016; indicated with *).

### C. Performance Improved and Feedback Adapted

We tested Hypothesis 3 with the **handedness** experiment (Fig. 6 and Fig. 7) to determine whether task performance changed with practice using time-domain reference tracking error ||*r* − *y*||^2^ from (8). We found that performance improved rapidly within the first five trials and then did not change significantly, even after switching hands, regardless of which hand was used first (Fig. 7a). Performance improved significantly between the first and last five trials with the first hand (trials #1–5 and #26–30; Group RL: *Z* = 0.00, *p* = 0.004; Group LR: *Z* = 0.00, *p* = 0.004), and did not change significantly between the last five trials with the first hand and the first five trials of the second hand (trials #26–30 and #31–35; Group RL: *Z* = 21.0, *p* = 0.86; Group LR: *Z* = 19.0, *p* = 0.68). We did not find statistically significant differences between Group RL and Group LR in the first or last five trials with either hand (Mann-Whitney *U* : *p* > 0.05).

**Fig. 6:**
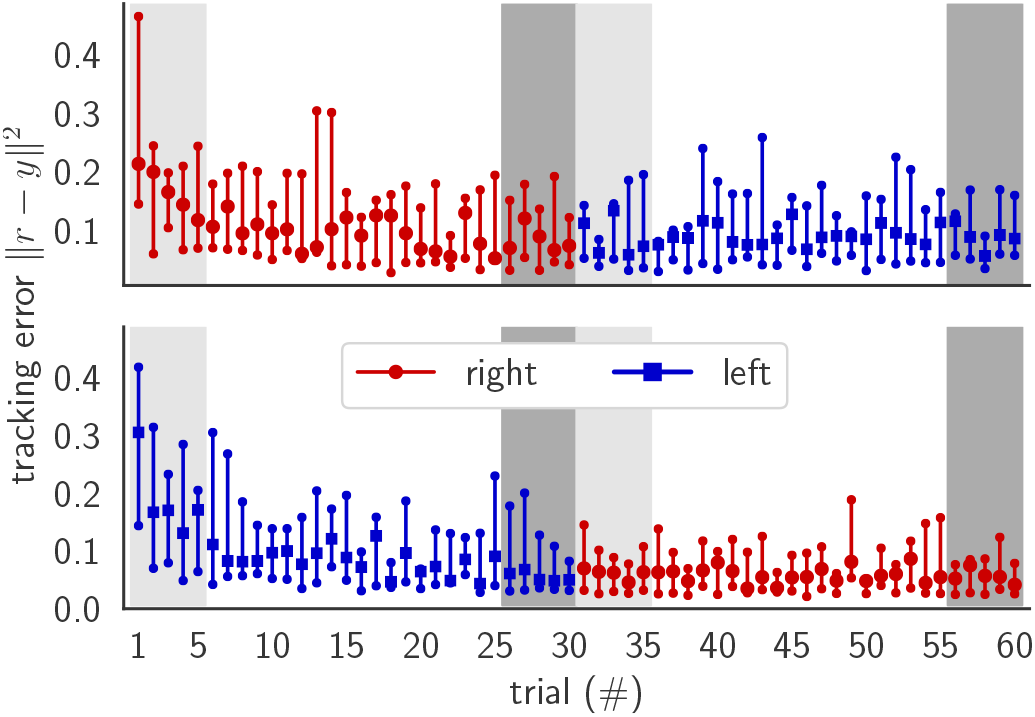
Tracking error from **handedness** experiment. Distributions (median, interquartile) of time-domain tracking error ||*r* − *y*||^2^ for 60 trials, with a switch between dominant (right; red circles) and non-dominant (left; blue squares) hands after trial 30, for two groups of 9 participants: (*top*) right then left (Group RL); (*bottom*) left then right (Group LR). Summary statistics in Fig. 7 use data from first five and last five trials with each hand, highlighted with light and dark gray boxes.

**Fig. 7:**
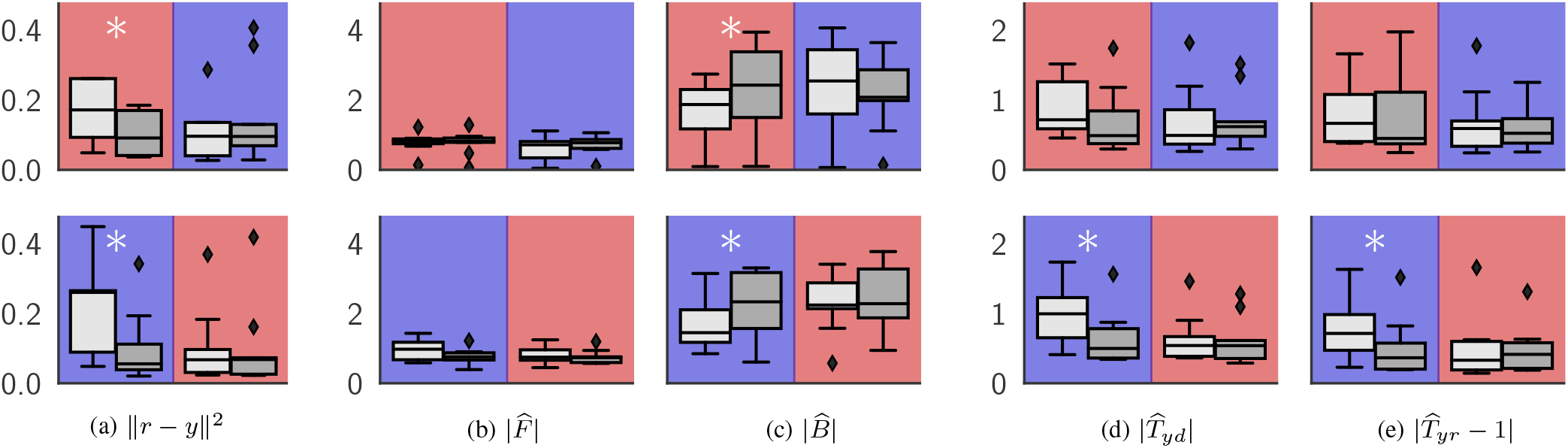
Summary statistics from **handedness** experiment. Distributions (median, interquartile, range) from first five (light gray box) and last five (dark gray box) of 30 trials with dominant (red solid background) and non-dominant (blue hatched background) hands: (a) tracking error ||*r* − *y*||^2^; mean magnitude of (b) feedforward 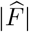 and (c) feedback 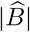 controllers (shared *y* axis); mean magnitude of (d) disturbance rejection 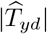 and (e) reference tracking 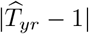 errors (shared *y* axis). Statistically significant (Wilcoxon signed-rank test: *p* < 0.05) differences between adjacent distributions indicated with*. Group RL in top row, Group LR in bottom row, as in Fig. 6.

To determine whether improvements in ||*r* − *y*||^2^ could be attributed to changes in feedback or feedforward control, we assessed whether feedback *B* or feedforward *F* control changed with practice using the mean magnitude of the frequency-domain representation 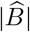 or 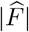 from (9). The mean magnitude of the feedback controller increased with practice for both groups (*Z* = 3.0, *p* = 0.02 in both) between the first and last five trials with the first hand, and did not change when switching to the second hand (*p* > 0.05). There was no statistically significant change in the mean magnitude of the feedforward controller across all conditions (*p* > 0.05) (Fig. 8). We did not find statistically significant changes in the phase of *F* and *B* at any stimulated frequency.

**Fig. 8:**
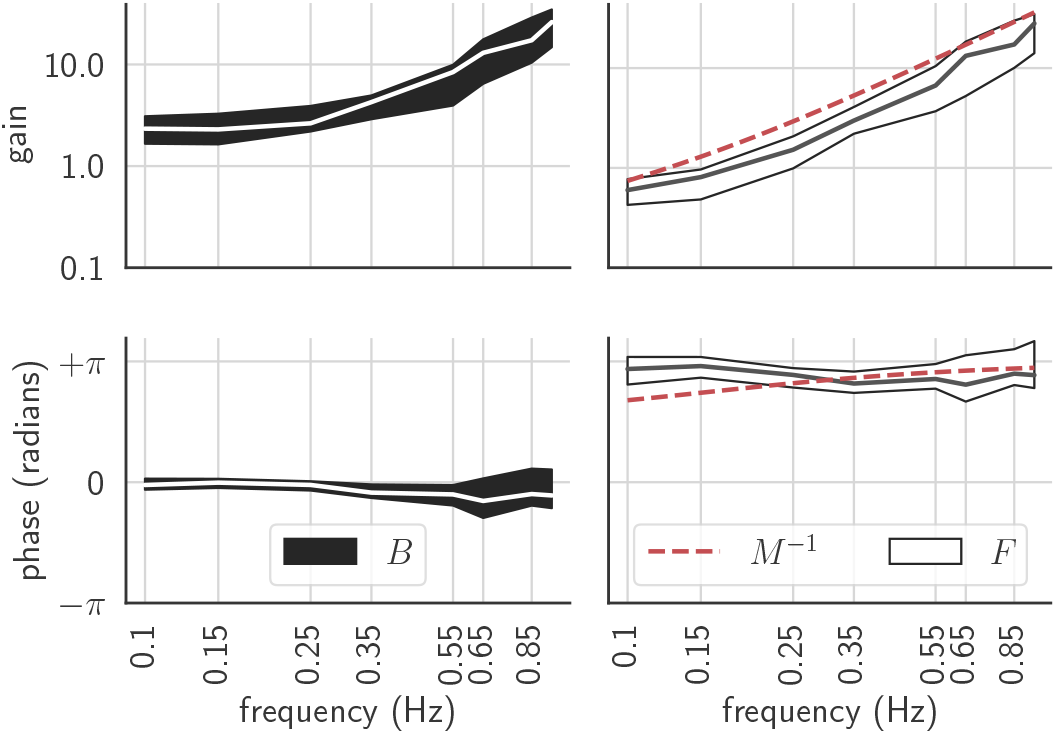
Human feedback (B) and feedforward (F) controllers. Distributions (median, interquartile) obtained by pooling data from the last five trials with each hand for both groups in the **handedness** experiment; we did not observe statistically significant differences between groups or hands (Wilcoxon signed-rank test: *p* > 0.05).

We observed system-level performance improvements at the first two stimulated frequencies (0.10, 0.15 Hz) solely for Group LR (Fig. 7). Group LR significantly decreased both 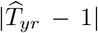 (*Z* = 4.0, *p* = 0.028) and 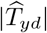 (*Z* = 0.0, *p* = 0.004) through experience with their first (left) hand, indicating significant improvements in reference tracking and disturbance rejection. This improved performance persisted even after switching from the left hand to the right hand, suggesting some transfer of knowledge between hands.

### D. User Response Diminished at Non-Stimulated Frequencies

Although we saw significant improvements in tracking performance with practice, we only observed modest or no improvements in system-level performance at stimulated frequencies. These results led us to consider user response at non-stimulated frequencies, since attenuating such response improves tracking performance. For both groups, the magnitude of the response at non-stimulated frequencies below crossover (0.25 Hz) decreased significantly between the first and last five trials with the first hand (trials #1–5 and #25–30) (Fig. 9), and this diminished response transferred between hands.

**Fig. 9:**
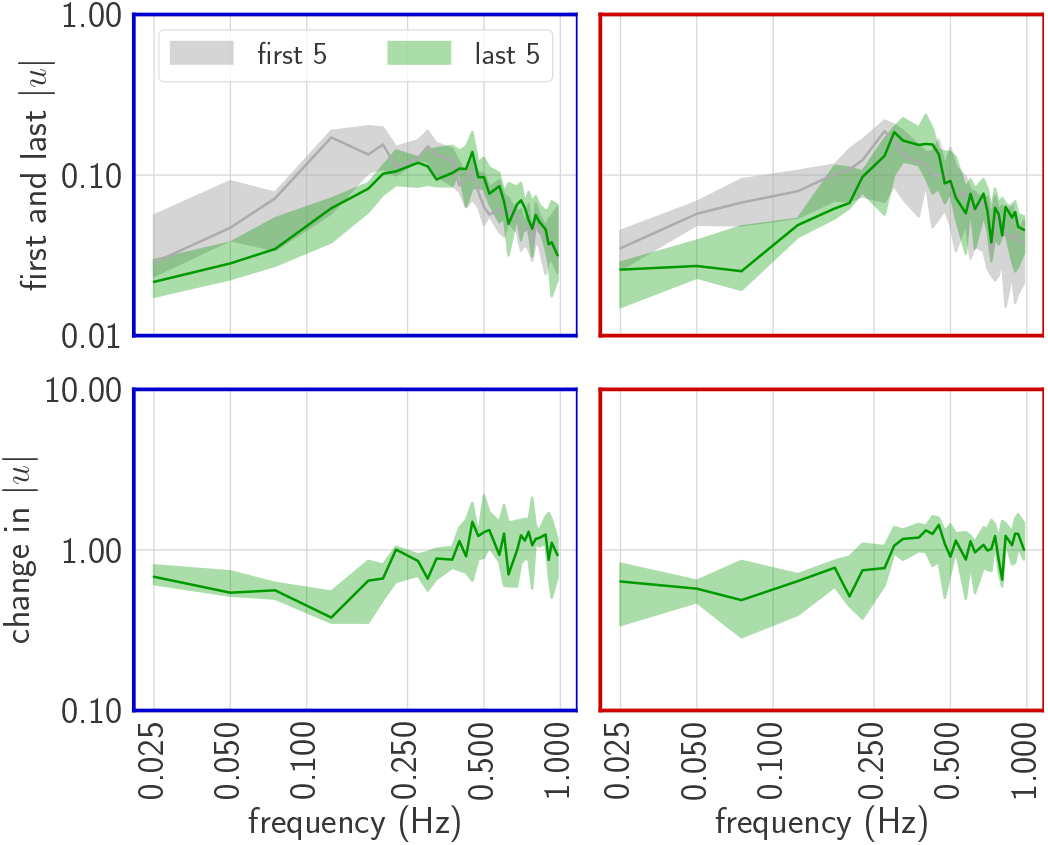
Change in effect of sensorimotor noise. (*top row:*) Distributions (median, interquartile) of magnitude of user response at non-stimulated frequencies from first and last five trials with first hand (trials #1–5 in light gray and #26–30 in green) in **handedness** experiment. (*bottom row:*) Ratio of user response magnitudes between first and last five trials with first hand decreases significantly below crossover (0.25 Hz). Group LR in left column, Group RL in right column.

## V. Discussion

Prior work demonstrated that people adapt feedback and feedforward controllers differently with the dominant and non-dominant hands during reaching tasks [13], [15]–[17]. However, little is known about how handedness affects learned controllers in continuous trajectory-tracking tasks such as the one considered in this study. When subjects reach to targets, feedback and feedforward control are assumed to be episodic: the initial ballistic motion is attributed to solely feedforward control (since sensorimotor delays preclude feedback) whereas corrective motions in the latter stage of the reach are attributed to solely feedback control [13], [15]–[17]. In contrast, feedback and feedforward processes are engaged simultaneously when subjects track continuous trajectories as in our experiments.

To assess how feedback and feedforward controllers are learned through experience and transferred between hands in a trajectory-tracking task, we extended, validated, and applied a non-parametric system identification method (adapted from [2], [5], [6], [8]). We found that feedback and feed-forward controllers estimated for different hands were not distinguishable and that learned controllers transferred between hands. Trajectory-tracking performance improved significantly with practice, but system-level performance improvements were significant only for the group that learned the trajectory-tracking task with their non-dominant hand first. Surprisingly, we did not find significant adaptation of the feedforward controller across the sample population. Instead, performance improvements can be attributed to a significant increase in feedback gain below crossover frequency; this accounts for significant changes in the effect of disturbances applied both externally by the experimenter and internally by sensorimotor noise.

### A. Response to Reference and Disturbance (Approximately) Superimposed Across Conditions

We found small but statistically significant differences between the transformations *T_ud_, T_ur_* estimated using data from disturbance-only (0, *d*) and reference-only (*r,* 0) trials and the combined reference-and-disturbance (*r, d*) trials. Thus, the controllers implemented by our participants to control a second-order system do not satisfy the superposition principle (1) as well as in our previous findings for first-order systems [5]. We attribute this difference to the increased difficulty of the trajectory-tracking task for a second-order system. However, considering how different each of the experimental conditions in Fig. 3 are from the user’s perspective – namely, that (*r,* 0) trials only have reference, (0, *d*) trials only have disturbance, and (*r, d*) trials have both reference and disturbance – we regard the empirical transformations in Fig. 4 as remarkably consistent across the qualitatively different conditions in Fig. 3.

Similarly to our previous findings for first-order systems [5], we found higher variability in estimates of transformation magnitude at higher frequencies compared to lower frequencies. Thus, although we found evidence that our human-in-the-loop control system is mildly nonlinear, neglecting this nonlinearity nevertheless yields good predictions for the human’s learned controllers, so our results support Hypothesis 1 with caveats.

Although human behavior is richly varied and nonlinear in general, our results support the assumption that people can behave remarkably linearly after sufficient experience interacting in closed-loop with a linear time-invariant system [1], [5]–[8], [32], [33]. Previous studies have ensured that human-in-the-loop-systems are approximately linear by using experts such as pilots [2] or only collecting data after participants undergo practice [1], [33]. Because our experiments commenced immediately without providing time for participants to explore the interface or machine dynamics (let alone become experts), this lack of practice may have contributed to the mild nonlinearities we observed. Future studies may benefit from estimation of nonlinearity [32], especially during learning.

### B. Combined Feedback and Feedforward Improved Prediction

We observed significant improvements in prediction accuracy with the feedback-plus-feedforward model compared to the feedback-only model at all frequencies in Fig. 5. This improvement in prediction accuracy implies that a model selection procedure based on an information criterion [34] would favor the combined feedback-plus-feedforward model over the feedback-only model if prediction accuracy was prioritized over model simplicity. Thus, our results lend further support for Hypothesis 2, consistent with previous results for first-order [1], [5], [6], [8] and fourth-order [7], [33] systems.

Our system identification method assumes the human controller consists of parallel feedback and feedforward controllers. However, the method does not assume or require either controller to be non-zero; in particular, if participants did not employ feedforward control, our method would yield a feedforward estimate with negligible magnitude. We emphasize that including *both* reference-tracking *and* disturbance-rejection in the task is necessary to ensure we can solve two independent equations in two unknowns (6) to uniquely determine feedback and feedforward controllers using our non-parametric modeling method.

### C. Performance Improved Because Feedback Adapted

Regardless of which hand was used first, participants significantly improved tracking performance through experience with their first hand. This improvement in time-domain performance persisted when participants switched hands, suggesting that learned controllers transferred between hands. Since we observed corresponding significant increases in feedback gain and observed no significant change in feedforward, we attribute this performance improvement to changes in feedback. These findings lead us to *reject* Hypothesis 3.

Our Hypothesis 3 was motivated by previous studies of human sensorimotor learning during reaching tasks that suggest improvements in end-point precision were due to improvements in initial movement (feedforward control) for the dominant (right) hand and improvements in error correction (feedback control) for the non-dominant (left) hand [13], [15]– [17]. However, there are differences between target-reaching tasks and the trajectory-tracking task used in this current experiment. For instance, the target-reaching tasks in [13], [15]–[17] are brief (approximately 1 sec in duration), so feedforward control is thought to dominate user response for a significant fraction of each trial since visual feedback is delayed by approximately 250 msec, and the target’s location changes discontinuously when the trial begins. In contrast, feedback and feedforward are engaged simultaneously for the entire 40 sec duration of each of our trajectory-tracking trials, and the reference changes continuously throughout the trial. The differences in experiment design could account for the differences we observed in how feedback and feedforward adapt. Since increasing the difficulty of a target-reaching task affects adaptation of feedback and feedforward [35], [36], it is possible that changing the machine dynamics or user interface may affect adaptation of feedback and feedforward in trajectory-tracking tasks.

Our inability to detect adaptation in feedforward control over a 1-hour period is inconsistent with previously published research that demonstrated adaptation of feedforward control over a 2-week period [7]. However, there are significant differences between our study methodology and [7] that may explain why we did not observe feedforward adaptation. First, the participants in [7] were tasked with learning to track a fourth-order system, which is significantly more complex than the second-order system used here, and the differing location of machine poles and zeros may affect learned controllers and tracking performance [37]. Second, since many of our participants reported prior experience controlling second-order systems (e.g. driving cars, playing video games), they may have employed a previously-learned feedforward controller in our experiment. Third, there was a significant difference in practice time between the two studies. In [7], participants learned the system dynamics over two weeks, whereas in our study, participants learned the system dynamics over 1 hour. Although we observed performance plateau during the 1-hour study, a longer practice time over the course of days or weeks may result in significant adaptation of feedforward control. Finally, and most significantly, while participants in [7] were tasked with following a predictable chirp trajectory, we tasked our participants to track unpredictable sum-of-sine trajectories. Stimuli predictability is known to affect tracking performance for human-in-the-loop systems [6, Fig. 5] [38, Fig. 5], possibly due to the use of internal *signal generators* [39], [40] (as opposed to the internal *controllers* posited here).

### D. Adaptation of Feedback Improved System-Level Performance For Group LR

To determine whether adaptations in feedback controller gain lead to system-level improvements in performance, we looked for differences in 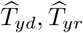 at the first two stimulated frequencies (0.10, 0.15 Hz) by comparing the first five and last five trials with each hand. For Group LR, we saw improvements in both 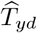 and 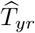 with their first hand, suggesting that reference tracking and disturbance rejection both improved. Despite clear improvements in time-domain performance for both groups, we did not observe statistically significant improvements in system-level performance at the stimulated frequencies for Group RL.

One possible explanation for these findings is that participants may initially find it more challenging to perform the trajectory-tracking task with their non-dominant hand, producing a larger effect that was easier to detect statistically. Consideration of sample size provides an alternative explanation for observed system-level differences in group performance that points to interesting directions for future study. Group LR and Group RL were relatively small populations (9 participants in each group), so there may have been unmeasured group-level differences. For instance, participants reported subjective differences in the strategy they employed to improve tracking performance. Some participants acknowledged that they were controlling the cursor acceleration and consciously altered their response accordingly, while others mainly focused on reactively minimizing tracking error. Future experiments with a larger number of participants are needed to determine whether different subpopulations employ different strategies when learning controllers.

### E. Adaptation of Feedback Affected the Effect (but not the Source) of Sensorimotor Noise

Since time-domain tracking performance improved significantly for both groups of participants but rejection of disturbance stimuli and tracking of reference stimuli only improved for one group, we are led to consider user response at frequencies we measured but did not stimulate. Any user response at non-stimulated frequencies degrades time-domain tracking performance, so it is in the users’ best interest to suppress this response [41]. We observed nonzero user response at non-stimulated frequencies, and this response decreased significantly with practice for frequencies below crossover for the first hand in both groups (Fig. 9). This suggests that, instead of (or in addition to) improving performance of disturbance rejection and trajectory-tracking at stimulated frequencies, the participants suppressed their response at low non-stimulated frequencies, leading to improved time-domain performance.

Because the machine dynamics and feedback in Fig. 1(b) are linear time-invariant, the user response at non-stimulated frequencies arises due to (i) nonlinearity in the human’s transformation and/or (ii) sensorimotor noise. Although we found evidence for (i) mild nonlinearities (see Fig. 4 and Sec. V-A), we tested for but did not find significant coherent responses in the user response at harmonics of the stimulated frequencies (i.e. non-stimulated frequencies), so nonlinearity alone does not appear to explain our observations. Assuming instead that user response at non-stimulated frequencies arises solely due to (ii) additive sensorimotor noise, we did not find statistically significant changes in this noise with experience. Indeed, despite the fact that we observed significant changes in feedback *B* and user response *u* at non-stimulated frequencies, we observed no significant changes in the power spectrum of the imputed disturbance *δ* = (1 + *MB*)*u*. Instead, the *effect* of the noise was attenuated by the increase in feedback gain below crossover. This result is consistent with prior studies from sensorimotor control that found the presence of significant noise whose statistics did not change with the limited amount of practice (less than 1 hour) considered here [42].

### F. Does Stimulus or Noise Drive Learning?

When learning to perform novel tasks like controlling a cursor on a screen or reaching under a force field, sensorimotor noise and movement variability are crucial for driving learning [43]–[45]. As people explore the action space for a particular task, certain movements (e.g. tracking a trajectory with specific frequency components) result in greater reward (e.g. improved tracking) [43]. With significant practice, noise and variability decreases, leading to improved performance in ballistic throwing [42], [46] and reaching [45] tasks. Similarly, we argue here that our observations that 1) there was time-domain improvement, 2) there was no corresponding system-level performance improvement at stimulated frequencies, and 3) user response decreased at non-stimulated frequencies below crossover, suggests that reducing the effect of sensorimotor noise may be a crucial aspect of performance improvement in continuous trajectory-tracking tasks. Although out of scope for our study, our results indicate that changes in sensorimotor noise at non-stimulated frequencies should be considered in addition to feedback and feedforward control at stimulated frequencies in studies of human-in-the-loop control systems.

## VI. Conclusion

Understanding how humans learn to track continuous trajectories with their dominant and non-dominant hands is crucial for enabling bimanual device control when teleoperating a surgical robot or manipulating objects in augmented or virtual reality. To this end, we first validated a non-parametric modeling method to simultaneously estimate feedback and feedforward control during a second-order continuous trajectory-tracking and disturbance-rejection task with seven participants. We then investigated adaptation of feedback and feedforward control and corresponding system-level changes in performance when nine participants learned to track with their right hand before their left hand, and when nine other participants learned to track with their left hand before their right hand.

Our study demonstrated that: (1) feedback control adapted with practice and transferred between hands in both groups; (2) feedback adaptation improved system-level performance in tracking prescribed references and rejecting externally-applied disturbances for the group that first learned the task with their non-dominant (left) hand; and (3) feedback adaptation improved tracking performance by attenuating the effect of a user’s sensorimotor noise in both groups. These findings suggest that handedness may not affect learned controllers, demonstrate that learned controllers may be transferred between hands, and highlight the importance of attenuating sensorimotor noise for human-in-the-loop control systems.

## Acknowledgments

We thank our participants for sharing their time with us, and we thank Pavan A. Vaswani and the anonymous reviewers for invaluable feedback provided on earlier versions of this manuscript. This research was funded by the University of Washington Institute for Neuroengineering (UWIN), the Lab for Amplifying Movement and Performance (AMP Lab) at the University of Washington, and the National Science Foundation Cyber-Physical Systems Program (Award #1836819).

## Appendix A Computing User Input Prediction

To evaluate Hypothesis 2, where we use data from the **linearity** experiment to compare the predictive accuracy of *B*-only and *B* + *F* models, we partitioned data from the conditions in TABLE II into disjoint *train* and *test* subsets as follows. First, we compute *B* at each stimulated frequency for each participant by averaging (6a) across the ten (0, *d*) trials. Subsequently, for the *B* + *F* model, we use each participant’s estimated *B* to compute their *F* at each stimulated frequency by averaging (6b) across the ten (*r,* 0) trials. Thus, the *train* dataset consisted of ten (0, *d*) trials and ten (*r,* 0) trials. We applied (5) to predict the (frequency-domain) user response for the *B* + *F* model in the last ten (*r, d*) trials using the user’s *B* and *F* estimates.

For the *B*-only model, we set *F* = 0 in (5) so that

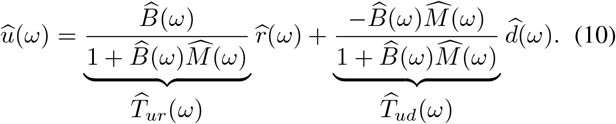

We can again compute *B* at each stimulated frequency for the ten (0, *d*) trials by averaging (6a). Then, we used (11) to compute *B* during the ten (*r,* 0) trials, and took the average to obtain an estimate of *B*.

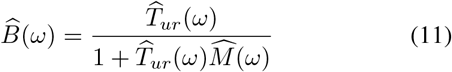

We applied (10) to predict user response for the *B*-only model in the last ten (*r, d*) trials using the user’s *B* estimates.

By performing this analysis at each stimulated frequency for each of the ten (*r, d*) trials for each participant, we obtained a predicted user response 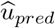 which we compared to the measured user input 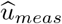. Thus, the *test* dataset consisted of ten (*r, d*) trials.

## Appendix B Computing *R*^2^ for User Input Prediction

We used the coefficient of determination *R*^2^ [31, Eqn. (3.9)] to assess prediction accuracy of the user input 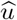 at each frequency. For each set of 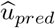 and 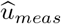 obtained from the (*r, d*) trials (see Appendix A), we computed an *R*^2^ value at each stimulated frequency for each trial *i* with the equation,

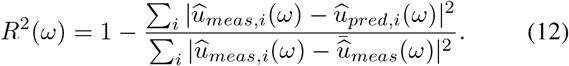

We defined 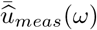 as the average user input of the measured input in the ten (*r, d*) trials that were used to make a prediction. We computed each of these quantities in the frequency domain using complex numbers, so we computed an *R*^2^ value at each stimulated frequency that represents deviation in both magnitude and phase. If the predictions perfectly match the measured user inputs, then *R*^2^ = 1. If the predictions do no better than predicting the mean of the observed user inputs, then *R*^2^ = 0. If the predictions are worse than predicting the mean of the observed user inputs, then *R*^2^ < 0.

## Appendix C Wilcoxon Signed-Rank Test

The Wilcoxon signed-rank test is a non-parametric paired *t*-test for data that is not normally distributed [30, Sec. 5.7]. Parametric statistical tests like the paired *t*-tests come with several assumptions that must be verified, such as that the data must be normally distributed [30]. We determined that the assumption of normality does not hold for our dataset using the Shapiro-Wilk test (*α* = 0.05) [47]. Therefore, we chose to use the Wilcoxon signed-rank test.

The Wilcoxon signed-rank test compares whether the differences between two conditions for a single group of *N* individuals have statistically different medians or not. The test does this by ranking the absolute difference between the two conditions for each participant, with 1 being assigned to the individual with the smallest difference, and *N* being assigned to the individual with the largest difference. Then, the rank for each individual is multiplied by 1 or −1 depending on whether the difference between the two conditions were positive or negative. The test statistic *Z* is computed as the sum of the signed ranks, and the *p*-value can be defined from the computed *Z* value [30, Sec. 5.7].

Because the test compares differences between all samples from two datasets, it is generally not possible to determine whether there is a statistically significant difference between the two datasets using only the median and interquartile statistics represented in box plots.

## Footnotes

1 Frequency where gain of loop transfer function 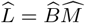 equals 1 [2].

2 Demographics were not recorded for the **linearity** experiment.

## References

[1] F. M. Drop, D. M. Pool, H. J. Damveld, M. M. van Paassen, and M. Mulder, “Identification of the feedforward component in manual control with predictable target signals,” IEEE Transactions on Cybernetics, vol. 43, no. 6, pp. 1936–1949, 2013.

[2] D. T. McRuer and H. R. Jex, “A review of quasi-linear pilot models,” IEEE Transactions on Human Factors in Electronics, no. 3, pp. 231–249, 1967.

[3] E. Roth, D. Howell, C. Beckwith, and S. A. Burden, “Toward experimental validation of a model for human sensorimotor learning and control in teleoperation,” in SPIE Conference on Micro-and Nanotechnology Sensors, Systems, and Applications IX, vol. 10194, 2017.

[4] K. van der El, D. M. Pool, M. R. M. van Paassen, and M. Mulder, “Effects of preview on human control behavior in tracking tasks with various controlled elements,” IEEE Transactions on Cybernetics, vol. 48, no. 4, pp. 1242–1252, 2017.

[5] M. Yamagami, D. Howell, E. Roth, and S. A. Burden, “Contributions of feedforward and feedback control in a manual trajectory-tracking task,” in IFAC Conference on Cyber-Physical-Human Systems (CPHS), vol. 51, no. 34. Elsevier, 2018, pp. 61–66.

[6] B. Yu, R. B. Gillespie, J. S. Freudenberg, and J. A. Cook, “Human control strategies in pursuit tracking with a disturbance input,” in IEEE Conference on Decision and Control (CDC), 2014, pp. 3795–3800.

[7] X. Zhang, S. Wang, J. B. Hoagg, and T. M. Seigler, “The roles of feedback and feedforward as humans learn to control unknown dynamic systems,” IEEE Transactions on Cybernetics, vol. 48, no. 2, pp. 543–555, 2017.

[8] M. Yamagami, K. M. Steele, and S. A. Burden, “Decoding intent with control theory: comparing muscle versus manual interface performance,” in ACM Conference on Human Factors in Computing Systems (CHI), 2020, pp. 1–12.

[9] M. Mulder, D. M. Pool, D. A. Abbink, E. R. Boer, P. M. Zaal, F. M. Drop, K. van der El, and M. M. van Paassen, “Manual control cybernetics: state-of-the-art and current trends,” IEEE Transactions on Human-Machine Systems, vol. 48, no. 5, pp. 468–485, 2017.

[10] K. Flowers, “Handedness and controlled movement,” British Journal of Psychology, vol. 66, no. 1, pp. 39–52, 1975.

[11] R. C. Oldfield, “The assessment and analysis of handedness: the Edinburgh inventory,” Neuropsychologia, vol. 9, no. 1, pp. 97–113, 1971.

[12] D. Kilshaw and M. Annett, “Right-and left-hand skill I: Effects of age, sex and hand preference showing superior skill in left-handers,” British Journal of Psychology, vol. 74, no. 2, pp. 253–268, 1983.

[13] S. V. Duff and R. L. Sainburg, “Lateralization of motor adaptation reveals independence in control of trajectory and steady-state position,” Experimental Brain Research, vol. 179, no. 4, pp. 551–561, 2007.

[14] R. S. Johansson, A. Theorin, G. Westling, M. Andersson, Y. Ohki, and L. Nyberg, “How a lateralized brain supports symmetrical bimanual tasks,” PLoS Biology, vol. 4, no. 6, p. e158, 2006.

[15] R. L. Sainburg and D. Kalakanis, “Differences in control of limb dynamics during dominant and nondominant arm reaching,” Journal of Neurophysiology, vol. 83, no. 5, pp. 2661–2675, 2000.

[16] L. B. Bagesteiro and R. L. Sainburg, “Handedness: dominant arm advantages in control of limb dynamics,” Journal of Neurophysiology, vol. 88, no. 5, pp. 2408–2421, 2002.

[17] C. N. Schabowsky, J. M. Hidler, and P. S. Lum, “Greater reliance on impedance control in the nondominant arm compared with the dominant arm when adapting to a novel dynamic environment,” Experimental Brain Research, vol. 182, no. 4, pp. 567–577, 2007.

[18] J. A. Anguera, C. A. Russell, D. C. Noll, and R. D. Seidler, “Neural correlates associated with intermanual transfer of sensorimotor adaptation,” Brain Research, vol. 1185, pp. 136–151, 2007.

[19] K. van der El, S. Padmos, D. M. Pool, M. M. van Paassen, and M. Mulder, “Effects of preview time in manual tracking tasks,” IEEE Transactions on Human-Machine Systems, vol. 48, no. 5, pp. 486–495, 2018.

[20] K. J. Aström and R. M. Murray, Feedback systems: an introduction for scientists and engineers. Princeton university press, 2010.

[21] M. Olivari, F. M. Nieuwenhuizen, J. Venrooij, H. H. Bülthoff, and L. Pollini, “Methods for multiloop identification of visual and neuro-muscular pilot responses,” IEEE Transactions on Cybernetics, vol. 45, no. 12, pp. 2780–2791, 2015.

[22] C. L. Phillips, J. M. Parr, and E. A. Riskin, Signals, systems, and transforms. Prentice Hall, 2003.

[23] F. M. Drop, D. M. Pool, M. R. M. van Paassen, M. Mulder, and H. H. Bülthoff, “Objective model selection for identifying the human feedforward response in manual control,” IEEE Transactions on Cybernetics, vol. 48, no. 1, pp. 2–15, 2016.

[24] F. M. Drop, D. M. Pool, M. M. van Paassen, M. Mulder, and H. H. Bülthoff, “Effects of target signal shape and system dynamics on feedforward in manual control,” IEEE Transactions on Cybernetics, vol. 49, no. 3, pp. 768–780, 2018.

[25] K. van der El, D. M. Pool, H. J. Damveld, M. R. M. van Paassen, and M. Mulder, “An empirical human controller model for preview tracking tasks,” IEEE Transactions on Cybernetics, vol. 46, no. 11, pp. 2609–2621, 2015.

[26] M. A. Smith and R. Shadmehr, “Intact ability to learn internal models of arm dynamics in Huntington’s disease but not cerebellar degeneration,” Journal of Neurophysiology, vol. 93, no. 5, pp. 2809–2821, 2005.

[27] A. M. Zimmet, D. Cao, A. J. Bastian, and N. J. Cowan, “Cerebellar patients have intact feedback control that can be leveraged to improve reaching,” eLife, vol. 9, Oct. 2020.

[28] E. Roth, K. Zhuang, S. A. Stamper, E. S. Fortune, and N. J. Cowan, “Stimulus predictability mediates a switch in locomotor smooth pursuit performance for Eigenmannia virescens,” Journal of Experimental Biology, vol. 214, no. 7, pp. 1170–1180, 2011.

[29] F. M. Nieuwenhuizen, P. M. T. Zaal, M. Mulder, M. M. Van Paassen, and J. A. Mulder, “Modeling human multichannel perception and control using linear Time-Invariant models,” Journal of Guidance, Control, and Dynamics, vol. 31, no. 4, pp. 999–1013, Jul. 2008.

[30] W. J. Conover, Practical nonparametric statistics. John Wiley & Sons, 1999.

[31] S. A. Glantz, B. K. Slinker, and T. B. Neilands, Primer of Applied Regression & Analysis of Variance, Third Edition. McGraw-Hill Education, Apr. 2016.

[32] S. E. Wagenaar, D. M. Pool, H. J. Damveld, M. M. van Paassen, and M. Mulder, “Estimation of nonlinear contributions in human controller frequency response functions,” in IEEE Conference on Systems, Man, and Cybernetics (SMC), 2018, pp. 3434–3439.

[33] R. B. Warrier and S. Devasia, “Inferring intent for novice human-in-the-loop iterative learning control,” IEEE Transactions on Control Systems Technology, vol. 25, no. 5, pp. 1698–1710, 2016.

[34] N. M. Mangan, J. N. Kutz, S. L. Brunton, and J. L. Proctor, “Model selection for dynamical systems via sparse regression and information criteria,” Proceedings of the Royal Society A: Mathematical, Physical, and Engineering Sciences, vol. 473, no. 2204, p. 20170009, Aug. 2017.

[35] R. Seidler, D. Noll, and G. Thiers, “Feedforward and feedback processes in motor control,” Neuroimage, vol. 22, no. 4, pp. 1775–1783, 2004.

[36] C. J. Winstein, S. T. Grafton, and P. S. Pohl, “Motor task difficulty and brain activity: investigation of goal-directed reciprocal aiming using positron emission tomography,” Journal of Neurophysiology, vol. 77, no. 3, pp. 1581–1594, 1997.

[37] X. Zhang, T. M. Seigler, and J. B. Hoagg, “The impact of nonminimum-phase zeros on human-in-the-loop control systems,” IEEE Transactions on Cybernetics, 2020.

[38] S. A. S. Mousavi, F. Matveeva, X. Zhang, T. M. Seigler, and J. B. Hoagg, “The impact of command-following task on human-in-the-loop control behavior,” IEEE Transactions on Cybernetics, 2020.

[39] R. B. Gillespie, A. H. Ghasemi, and J. Freudenberg, “Human motor control and the internal model principle,” in IFAC Symposium on Analysis, Design, and Evaluation of Human-Machine Systems (HMS), vol. 49, 2016, pp. 114–119.

[40] S. Cutlip, J. Freudenberg, N. Cowan, and R. B. Gillespie, “Haptic feedback and the internal model principle,” in IEEE World Haptics Conference (WHC), Jul. 2019, pp. 568–573.

[41] J. Venrooij, M. Mulder, D. A. Abbink, M. M. Van Paassen, F. C. Van Der Helm, H. H. Bülthoff, and M. Mulder, “A new view on biodynamic feedthrough analysis: Unifying the effects on forces and positions,” IEEE Transactions on Cybernetics, vol. 43, no. 1, pp. 129–142, 2012.

[42] M. E. Huber, N. Kuznetsov, and D. Sternad, “Persistence of reduced neuromotor noise in long-term motor skill learning,” Journal of Neurophysiology, vol. 116, no. 6, pp. 2922–2935, 2016.

[43] A. K. Dhawale, M. A. Smith, and B. P. Ölveczky, “The role of variability in motor learning,” Annual Review of Neuroscience, vol. 40, pp. 479–498, 2017.

[44] D. J. Herzfeld and R. Shadmehr, “Motor variability is not noise, but grist for the learning mill,” Nature Neuroscience, vol. 17, no. 2, pp. 149–150, 2014.

[45] H. G. Wu, Y. R. Miyamoto, L. N. G. Castro, B. P. Ölveczky, and M. A. Smith, “Temporal structure of motor variability is dynamically regulated and predicts motor learning ability,” Nature Neuroscience, vol. 17, no. 2, pp. 312–321, 2014.

[46] C. J. Hasson, Z. Zhang, M. O. Abe, and D. Sternad, “Neuromotor noise is malleable by amplifying perceived errors,” PLoS Computational Biology, vol. 12, no. 8, p. e1005044, 2016.

[47] N. M. Razali, Y. B. Wah et al., “Power comparisons of Shapiro-Wilk, Kolmogorov-Smirnov, Lilliefors and Anderson-Darling tests,” Journal of Statistical Modeling and Analytics, vol. 2, no. 1, pp. 21–33, 2011.

